# Targeting mononuclear phagocytes for eradicating intracellular parasites

**DOI:** 10.1101/119297

**Authors:** Loris Rizzello, James D. Robertson, Philip M. Elks, Alessandro Poma, Nooshin Daneshpour, Tomasz K. Prajsnar, Dimitrios Evangelopoulos, Julio Ortiz Canseco, Simon Yona, Helen M. Marriott, David H. Dockrell, Simon J. Foster, Bruno De Geest, Stefaan De Koker, Timothy McHugh, Stephen A. Renshaw, Giuseppe Battaglia

## Abstract

Mononuclear phagocytes such as monocytes, tissue-specific macrophages and dendritic cells are primary actors in both innate and adaptive immunity, as well as tissue homoeostasis. They have key roles in a range of physiological and pathological processes, so any strategy targeting these cells will have wide-ranging impact. These phagocytes can be parasitized by intracellular bacteria, turning them from housekeepers to hiding places and favouring chronic and/or disseminated infection. One of the most infamous is the bacteria that cause tuberculosis, which is the most pandemic and one of the deadliest disease with one third of the world’s population infected, and 1.8 million deaths worldwide in 2015. Here we demonstrate the effective targeting and intracellular delivery of antibiotics to both circulating monocytes and resident macrophages, using pH sensitive nanoscopic polymersomes made of poly(2-(methacryloyloxy)ethyl phosphorylcholine)-co-poly(2-(diisopropylamino)ethyl methacrylate) (PMPC-PDPA). Polymersome selectivity to mononuclear phagocytes is demonstrated and ascribed to the polymerised phosphorylcholine motifs affinity toward scavenger receptors. Finally, we demonstrate the successful exploitation of this targeting for the effective eradication of intracellular bacteria that cause tuberculosis *Mycobacterium tuberculosis* as well as other intracellular parasites including the *Mycobacterium bovis, Mycobacterium marinum* and the most common bacteria associated with antibiotic resistance, the *Staphylococcus aureus*.

## Introduction

The human innate immune system - our frontline defence against potential pathogens - includes a range of effector cells. Examples are professional phagocytes, such as granulocytes (*i.e*., basophils, eosinophils and neutrophils) and mononuclear phagocytes (macrophages, dendritic cells, and monocytes).^1^ Phagocytes are responsible for the clearance of bacterial pathogens from the host, and have attracted much interest in the context of focused antimicrobial drug delivery. In parallel, some of the most deadly pathogens have acquired the ability to evade the phagocytes’s unique panel of molecular defences. As a consequence, while such phagocytes have evolved to eradicate invading pathogens, certain bacteria have simultaneously developed strategies to exploit macrophages as their preferential niche in which to evade host killing. This is known as the ‘macrophage paradox’ and it is the product of millions of years of co-evolution.^2^ Pathogens may inhabit different compartments in the macrophage; *Listeria monocytogenes, Shigella flexneri* and the *Rickettsiae rickettsii* proliferate within the macrophage cytosol, *Listeria pneumophila* colonises the ER-like vacuoles, and *Salmonella enterica* exploits the late endosomal compartments. More recently, a similar strategy has been reported for *Staphylococcus aureus*, suggesting that these bacteria are capable of hiding within professional phagocytes.^3^ The well studied intracellular pathogen, the *Mycobacterium tuberculosis*, survives within macrophages phagosomes, considered the most detrimental environment for pathogens. Yet, *M. tuberculosis* has evolved proteins that hinder phagosome maturation, and prevent its fusion with lysosomes.^4–6^ The optimal design of drug delivery systems should incorporate targeting specificity for the host cells type, the presence of the pathogen, and the pathogen sub-cellular location. Hence, there is a need for novel antibacterial compounds that combine potent antibacterial activity with the ability to cross biological barriers and finally reach the intracellular niche where the microorganisms hide - even more critical today with the emergence of drug resistant strains. Here we propose the use of synthetic vesicles - known as polymersomes - that are able to target infected phagocytes, to reach intracellular pathogens in their sub-cellular compartment, and locally release their antibacterial cargo. These polymersomes are formed through the self-assembly of amphiphilic copolymers in aqueous media,^7^ and combine the advantages of long-term stability with the potential to encapsulate a broad range of cargoes.^8–10^We have previously demonstrated that the pH sensitive block copolymer poly(2-(methacryloyloxy) ethyl-phosphorylcholine)- co-poly(2-(diisopropylamino)ethyl methacrylate) (PMPC-PDPA) can combine specific cellular targeting efficiency (through the PMPC block and its affinity toward the scavenger receptor B1),^11^ with effective endosomal escape and cytosolic delivery following internalisation (by the pH sensitive PDPA).^8,12^In this study, we describe the half life and bio-distribution of the PMPC-PDPA polymersomes *in vivo*, showing the dynamics of accumulation within different tissues. We then show the bio-molecular mechanism of cellular uptake and intracellular trafficking (namely, the polymersomes localisation in specific sub-cellular organelles). Finally, we demonstrate the potential of this strategy to revolutionise the treatment of several intracellular pathogens both *in vitro* and *in vivo*, namely *S. aureus*, M.bovis-attenuated Bacillus Calmette-Guérin (BCG), *M. tuberculosis*, and *M. marinum*. We demonstrate *in vitro* that PMPC-PDPA polymersomes loaded with antimicrobials (gentamicin, lysostaphin, vancomycin, rifampicin, and isoniazid) are able to decrease, and potentially even eradicate, these intracellular pathogens. Using embryos of the zebrafish (*Danio rerio*) as a model, we further demonstrate that encapsulated antimicrobials can effectively reduce the bacterial burden in disseminated infections *in vivo*.

## Results and discussion

PMPC-PDPA copolymers were synthesised using atom transfer radical polymerisation (ATRP) protocols and fully characterised by gel permeation chromatography (GPC) (Figure S1a) and NMR spectroscopy (Figure S1b). We used film hydration methods for the polymersome fabrication^13^ and we isolated monodisperse polymersomes from other structures by means of density gradient centrifugation.^14^ Transmission electron microscopy (TEM) and dynamic light scattering (DLS) analyses (**Figures S1c-d**) confirmed the purification processes were successful in isolating spherical polymersomes with homogeneous size and shape distributions. Finally, PMPC-PDPA was synthesised, bearing either Cy5 or Rhodamine dyes to facilitate imaging and quantification by fluorescence. We first studied the mechanisms of PMPC-PDPA polymersomes internalisation *in vitro* in macrophages using the monocytic cell line (THP-1).^15^ Live cell confocal laser scanning microscopy (CLSM) imaging of macrophages stained by CellMask™shows that the uptake of PMPC-PDPA polymersomes occurs rapidly (Figure 1a). We observed full saturation of the membrane within minutes of exposure, with several internalisation events within seconds of binding. The kinetics are shown in Figure 1b for four regions of interest (ROI) and all confirm rapid binding and endocytosis with very little difference between the different ROI, indicating a common uptake mechanism. We also CLSM-imaged and quantified the uptake of polymersomes at 8, 24 and 72 hours of incubation time (Figure 1c). Calcein (green) staining demonstrated that the cells remained viable for the incubation time tested. We further quantified the uptake using high performance liquid chromatography (HPLC) of the cell lysates after different incubation times. HPLC-based uptake quantifications revealed about 10^4^ polymersomes/cell after 8 hours, the number of polymersomes rose by 3 · 10^4^ after 24h and this remained constant up to 72 hours of incubation (Figure 1c). Even though a large number of polymersomes was internalised by the monocytes, viability assays (MTT) confirmed that free- and antimicrobial-encapsulated polymersomes do not affect the metabolic activity of THP-1 cells (Figure S2). We also performed quantitative PCR (qPCR) for a panel of genes, with the aim of identifying alterations of phagocyte homeostasis. We found that polymersomes did not affect the expression of the stress-related genes p21 and p53 (Figure 1e). In addition, incubation with polymersomes did not induce the oxidative stress genes CAT or SOD1, nor modify the cell metabolism genes CYP1Aq and CYP1B1. Neither did the polymersomes promote protein unfolding genes, as the sensors for the unfolded protein response (UPR) pathways, ATF4 and ATF6, were not increased by polymersomes treatment (Figure 1e).

**Figure 1:**
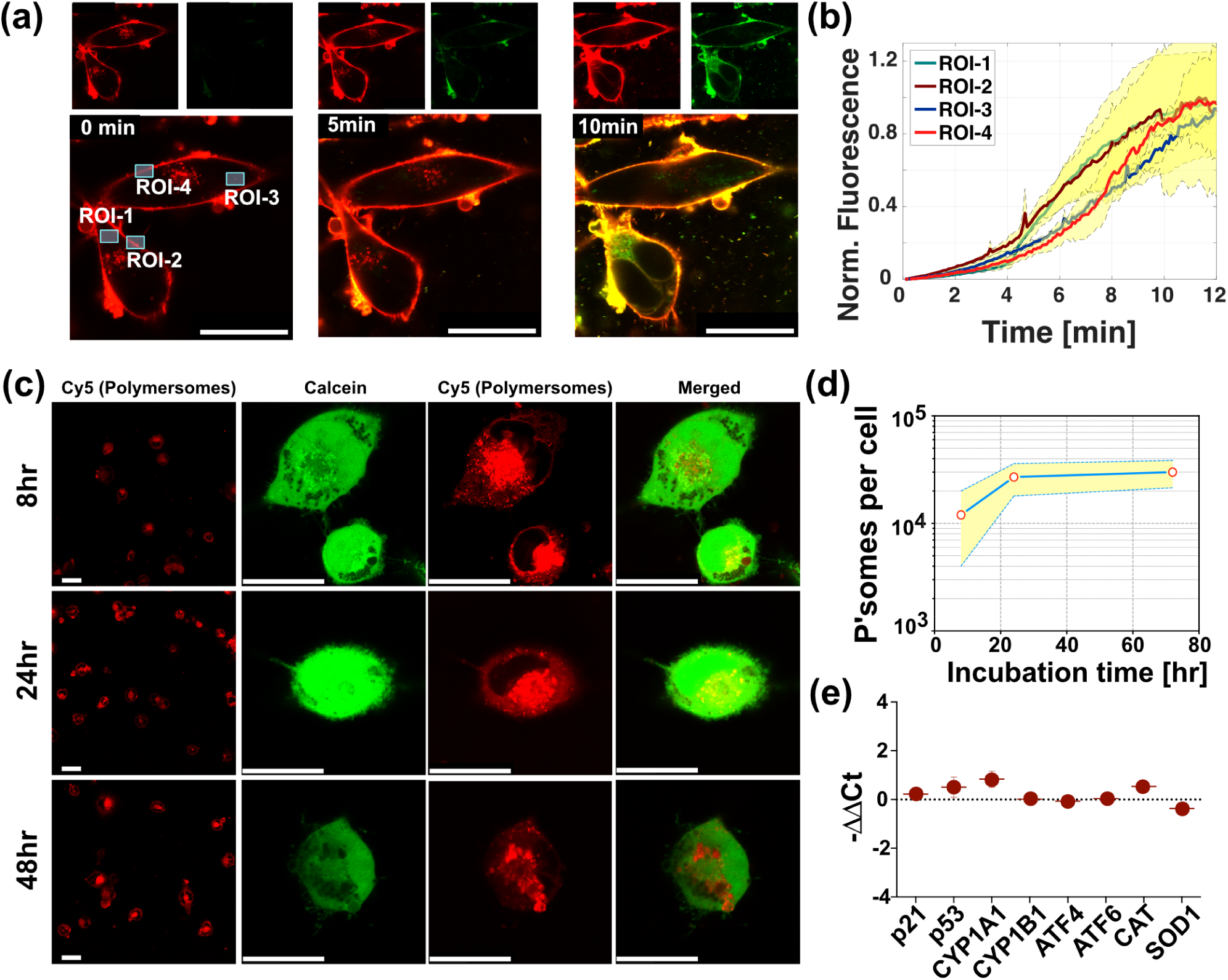
PMPC polymersome interact with phagocytes *in vitro*. **(a)** Real time imaging of polymersomes entering monocyte-derived macrophages (THP-1 cells) using confocal laser scanning microscopy (CLSM). Note the polymersomes (red signal) are labelled using Cy5, and the macrophage membrane (green signal) is stained using CellMask™. Scale bar = 25*μ*m. **(b)** Polymersome uptake measured in 4 different regions of interest (ROI) in (**a**) plotted as a function of time. **(c)** CLSM analyses of THP-1 cells incubated with rhodamine-labeled polymersomes for 8 hours, 24 hours, and 72 hours. Scale bar = 25*μ*m. **(d)** HPLC quantification of polymersome uptake into THP-1 cells at different incubation times. **(e)** Quantitative PCR (qPCR) for analysing the expression levels of genes involved in cell proliferation (p21 and p53), cell stress (CYP1A1 and CYP1B1), Unfolded Protein Response (ATF4 and ATF6), and oxidative stress (CAT and SOD).

In previous studies, we demonstrated that pH sensitive PMPC-PDPA polymersomes escape rapidly from early-endosomes.^12^ To test whether this was also the case in macrophages, Cy5-labelled polymersomes were incubated with THP-1 cells and a Cy3-LAMP1 lysosome marker. Cy5-polymersomes were found within macrophage lysosomes after 8 hours of incubation (Figure 2a). Similar results were observed after 24h and 72h of incubation with polymersomes (**Figures S3a-b**). These immunofluorescence experiments also confirmed that a considerable population of pH sensitive polymersomes escaped the endocytic pathways and moved to the cytosol (free red signal in macrophages), enabling access to the two intracellular compartments. This raised the question of whether the preferred driving force for polymersome internalisation is receptor-mediated endocytosis or phagocytosis. To investigate this, macrophages were incubated with the actin inhibitor cytochalasinB. We observed a complete inhibition of polymersome uptake, demonstrating that the entry process is mediated by actin-dependent transport (Figure 2b). Moreover, incubation with 15*μ*m and 30*μ*m dynasore (a dynamin inhibitor) reduced the polymersome uptake by 40% and 60%, respectively, but did not stop it completely (Figure 2b). This was unexpected, as the GTPase dynamin regulates membrane fission in clathrin-mediated endocytosis, as well as in phago- and macropinocytosis in eukaryotic cells.^16^ Few dynamin-independent entry pathways have been described, and they include the CDC42 (the preferred entry route of cholera toxin B),^17^ ARF1 and ARF6,^5, 18, 19^ and Flotillin 1 and 2 pathways.^20,21^ Our inhibition studies suggest that polymersomes can gain access through dynamin-independent endocytosis. Scavenger Receptors (SR) and SR-B1 in particular, are known receptors for PMPC-PDPA polymersome uptake in non-professional phagocytes.^11^ SR-B1 is known to play an critical role in pathogens recognition and in cholesterol homoeostasis.^22–24^ To test whether macrophages also internalise polymersomes using scavenger receptors, we incubated the cells with fucoidan, an inhibitor of Scavenger Receptors class A and B (SR-A and B). Despite the presence of the inhibitor, polymersomes were able to access macrophages, albeit with a considerable decrease in uptake of about 40% (Figure 2b). In order to define the contribution of the class A or B, macrophages were treated with polyinosinic acid (PA), a selective inhibitor of SR-A.^25^ Surprisingly, PA led to a significant increase in polymersome uptake (Figure 2b). This supports the involvment of SR-B1 - if SR-As were involved, the uptake would be hindered. Moreover, PA has been shown to bind to Tolllike receptor 3, inducing an increase in phagocytic activity,^25^ corresponding to an increase polymersome uptake. The concentrations of inhibitors used have been tested to study entry mechanisms while avoiding toxic effects to cells (Figure S4). Taken together, the inhibition-based studies indicate that polymersomes enter through active uptake, probably through a combination of dynamin-independent endocytosis and phagocytosis.

**Figure 2:**
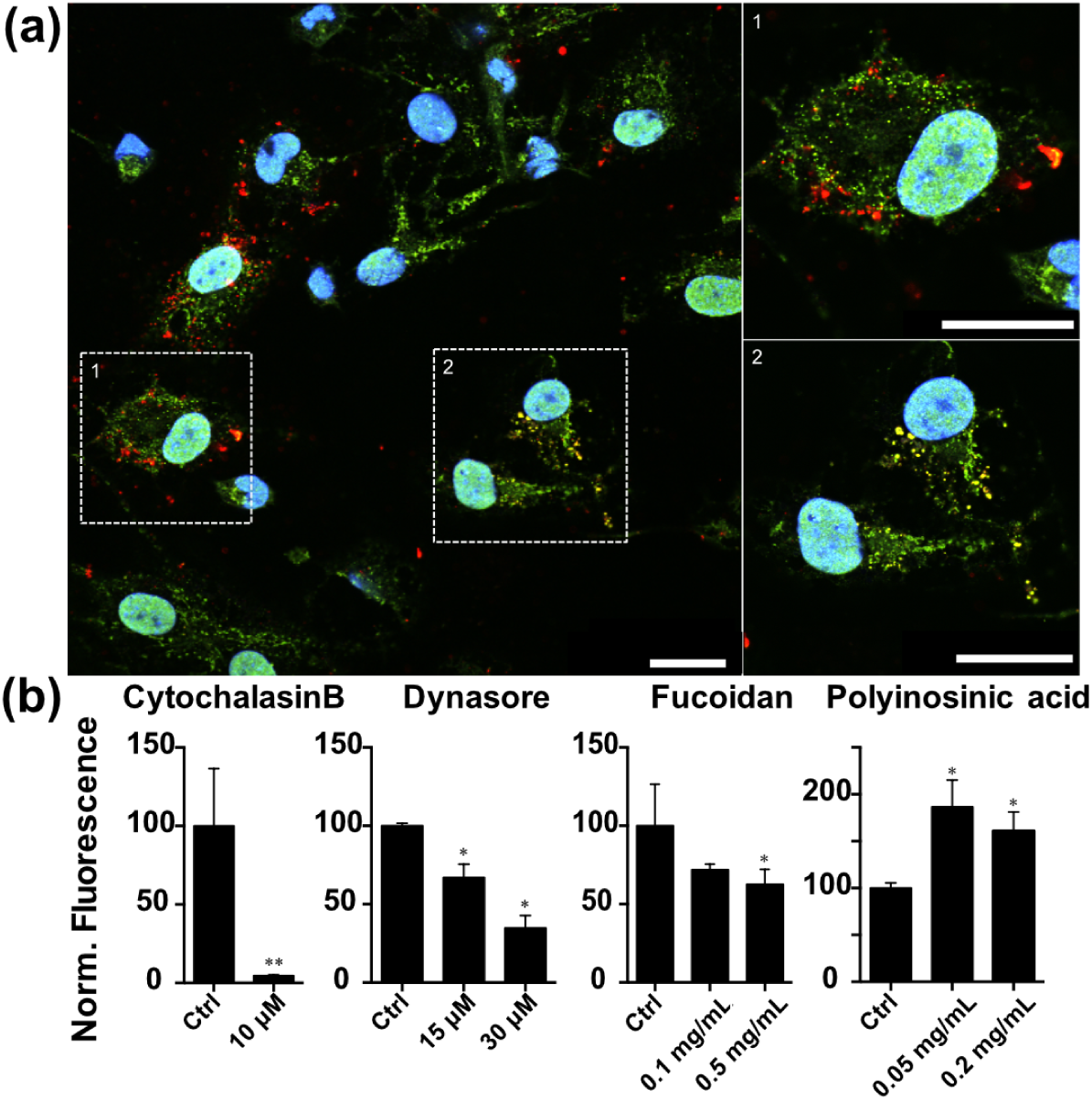
Macrophage internalisation of PMPC polymersomes mechanism is actin dependent transport. **(a)** CLSM investigation of THP-1 cells incubated with polymersomes for 8 hours. Polymersomes (Cy5-red) co-localise with LAMP-1 (green), indicating their presence in the lysosome (yellow merge). Scale bar = 25*μ*m. **(b)** Polymersome uptake after inhibition of different cellular components: CytochalasinB (actin inhibitor), Dynasore (dynamin inhibitor), Fucoidan (Scavenger Receptors A and B inhibitor), and Polyinosinic acid (Scavenger Receptor A inhibitor and Toll-like 3 receptor agonist ligand stimulator). (t-test compairason with *\*p* < 0.05).

The *in vitro* characterisation demonstrated that these polymersomes target and enter phagocytes through the scavenger receptors. We subsequently explored the inherent affinity of PMPC-PDPA polymersomes towards phagocytes *in vivo* using mice as model organism. Upon intravenous (i.v.) tail injection, we measured the plasma PMPC-PDPA *C_p_*(*t*) polymersomes concentration as a function of time (**Figure 3.a**). The plasma concentration decays according to two phases, 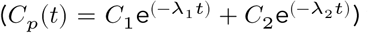 where about 84.5% of the polymersomes are quickly eliminated from the plasma with a fast half-life of 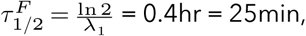 and the remaining amount is removed with slow half-life of 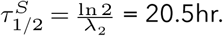 While these can be interpreted using compartmental pharmacokinetic models^26^ ascribing the biphasic decay to a combination of fast distribution and slower excretion, we opted here to factor the interaction with the blood cells, and the immune cells in particular, leaving more complex pharmacokinetic evaluation for future work. As a first assessment, we counted white and red blood cells at different times after i.v. polymersomes injections to check for any unwanted effects caused by polymersomes to blood cells. We did not detect any significant difference in cell counts (Figure S5). We thus measured the uptake of polymersomes in the different blood cell types by flow cytometry and observed a remarkable selectivity of PMPC-PDPA polymersomes for monocytes (Ly6C+ cells) with over 98% of these positive only 5min after i.v. injection (Figure 3b). This is in contrast to about 12% of B cells, 7% of T cells and 6% of neutrophils while red blood cells did not show any detectable level of polymersomes uptake. These are remarkable results considering monocytes make up only about 2% of the total blood cell population (Figure S5), strongly supporting the ability of PMPC-PDPA polymersomes to selectively target monocytes. The ability of polymersomes to distinguish sub-classes of monocytes including classical monocytes (Ly6C high) and non-classical monocytes (Ly6C low) was assessed. Classical monocytes internalised polymersomes more efficiently at early time points (after 10 minutes post injection, Figure 3c, red graph). However, 24 hours post injection, the non-classical monocytes more readily took up polymersomes (Figure 3c, blue graph). This shows we can target both types of monocytes, even though we can not exclude some of the polymersomes can be carried within monocytes during their transition from a classical (*i.e*., not having inflammatory attributes) to non-classical (pro-inflammatory) sub-set.^1^ These latter constitute 50% of the whole monocyte population (in mice). Such a strong interaction with monocytes suggests that polymersome distribution into other tissues can follow two routes, one direct from the blood and one piggy-backing in monocytes. To further assess this, organs excised at different time points were measured post injection and imaged after careful blood depletion by perfusion.^27^ There is a strong accumulation of polymersomes within the gastrointestinal (GI) tract and the liver (about 40% of the total fluorescence each)(Figure 3d). These large organs are followed by the spleen, kidneys, and lungs, while the other organs have the rest distributed among them. We can identify two regimes of distribution: (i) in the GI tract, liver, spleen, kidneys, lungs, and heart, the polymersome concentration peaks around the time (between 1 and 2 hrs) where 90% of the polymersomes are depleted from the blood plasma (**Figure. 3a**), (ii) in bone marrow, muscle, testis, thymus, spinal cord, and brain, the concentration, albeit quite low overall, rises steadily with time suggesting a very delayed distribution. The first group of organs has higher expression of SR-B1,^22–24^ and the data match the plasma circulation indeed, suggesting a fast diffusion from the blood to the tissues. In the other organs, particularly the poorly perfused, the steady rise in polymersome concentration could be explained by monocyte/macrophage penetration into tissues over time. In addition to the interaction between polymersomes and blood cells, we also studied the potential uptake by tissue resident macrophages. In particular, we chose to study the uptake in liver resident Kupffer cells, whose origin is mostly embryonic,^28^ and which have a critical role in cleaning the blood of pathogens and particulate materials.^29^ The selective uptake of polymersomes in liver resident macrophage Kupffer cells was seen in as little as 10min after i.v. injection (Figure 3e), and completely saturated after 24hr. Finally, we checked whether local phagocytes could be targeted by topical delivery, using intratracheal instillation to deliver rhodamine-labelled polymersomes directly to the lungs. The tissues were extracted 24h after administration and cells analysed by flow cytometry. We observed considerable targeting toward phagocytes, where over 90% of dendritic cells and macrophages in bronchoalveolar lavage, and 60% of macrophages and 15% of dendritic cells in the lung tissues, were positive for polymersome uptake.(Figure 3f)

**Figure 3:**
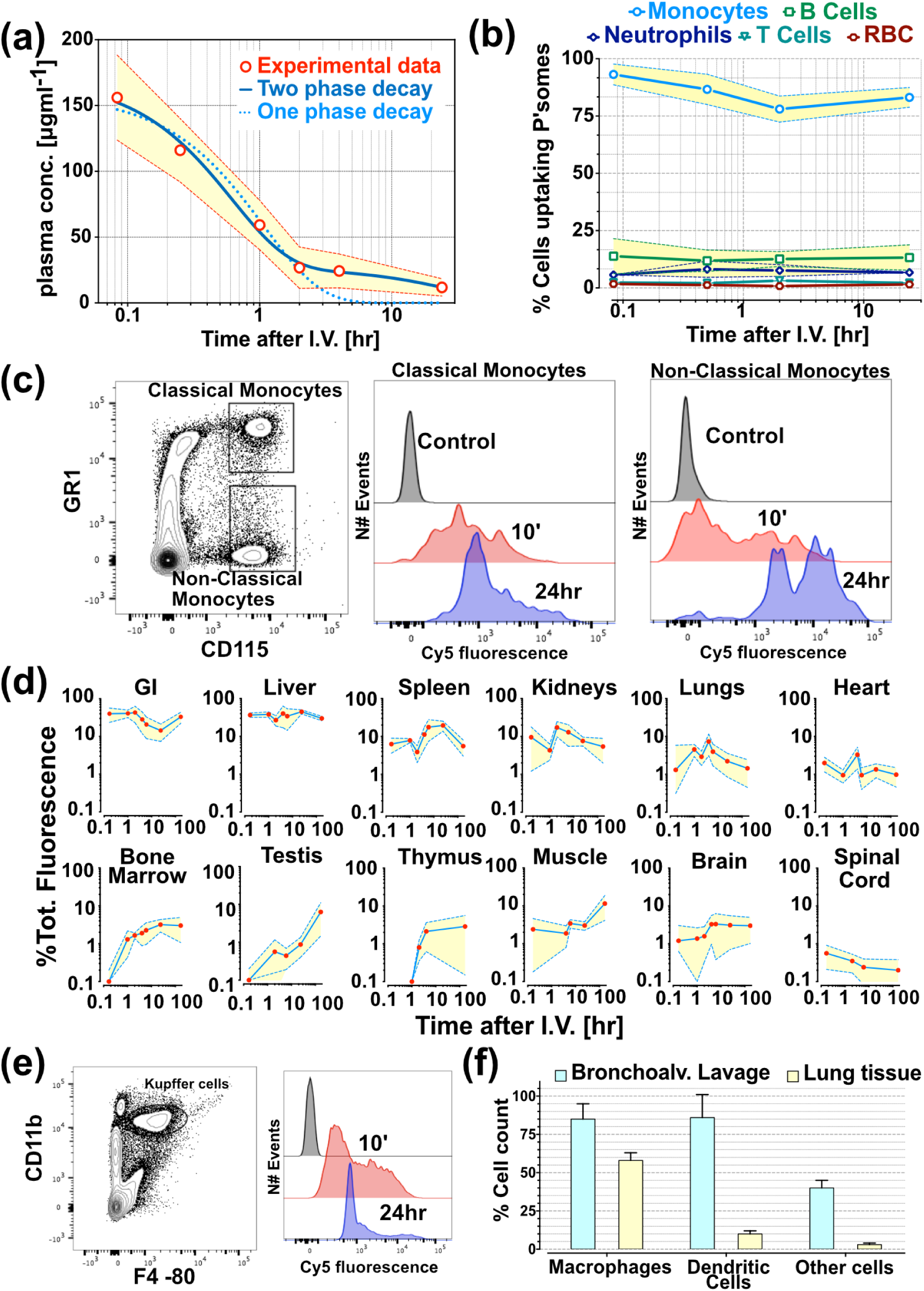
Bioavailability of polymersomes. **(a)** Plasma concentration of the PMPC-PDPA polymersomes as a function of time after i.v. injection. The data are fitted using a one-phase decay 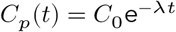 (dotted line) and a two-phase decay 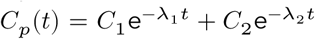 (solid line) corresponding to one- and two-compartment pharmacokinetic models, respectively. **(b)** Blood cells uptake of PMPC-PDPA polymersomes measured by flow cytometry. **(c)** Flow cytometry analysis of the interaction between PMPC-PDPA polymersomes and classical and non-classical monocytes. **(d)** *Ex vivo* analyses of PMPC-PDPA polymersome distribution in different organs at different times after i.v. injections. The data are showed as percentage of the total measured fluorescence (across all excised organs). **(e)** Flow cytometry analysis of the interaction between PMPC-PDPA polymersomes and liver resident macrophage kupffer cells. **(f)** Cell counts using markers for alveolar macrophage (M*ϕ*), dendritic cells (DC), and other cells, of both brochoalveoalar lavage and lung tissue 24hr after polymersome intratracheal instillation in mice.

Having demonstrated that PMPC-PDPA polymersomes are an optimal candidate for phagocyte targeting, their efficacy in delivering antimicrobials to reduce bacterial burden was examined using models of the intracellular pathogens *S. aureus, M.bovis-BCG, M. tuberculosis*, and *M. marinum*. The encapsulation efficiency of vancomycin, gentamicin, lysostaphin, rifampicin, isoniazid was assessed (Figure 4a), demonstrating our ability to load polymersomes with doses higher than the respective minimal inhibitory concentrations. It is important to note that the different drugs have considerable differences in molecular mass, hydrophilicity, and mechanism of action. Lysostaphin is a 27 KDa glycylglycine endopeptidase only soluble in water acting on the *S. aureus* cell walls. Gentamicin is a highly hydrophilic aminoglycoside that binds to the 30S subunit of the bacterial ribosome. Vancomycin is a relatively hydrophilic glycosylated nonribosomal peptide that inhibits cell wall synthesis. Rifampicin is a hydrophobic heterocyclic modified napthoquinone that inhibits bacterial DNA-dependent RNA synthesis. Finally, isoniazid is a small synthetic derivative of nicotinic acid with a poor water solubility that upon enzymatic activation inhibits the synthesis of mycoloic acids. These drugs are used clinically for the treatment of several infections and make a very diverse population of molecules to test the versatility of polymersomes. We thus tested the effect of the antimicrobials in the treatment of different infections by measuring the colony forming units (CFU) after increasing incubation periods. Treatment with polymersomes loaded with rifampicin or gentamicin improved the drug efficacy and reduced the number of viable *S. aureus* in THP-1 cells compared with controls (Figure 4b). Encapsulation of lysostaphin or vancomycin within polymersomes did not significantly improve or hinder drug efficacy. The enhancement of rifampicin and gentamicin at killing intracellular *S. aureus* compared to the same dose of free drugs can be ascribed to improved intracellular delivery and the consequent increase in drug reaching the intracellular pathogen. For both BTG and *M. tuberculosis*, we limited our screening to rifampicin and isoniazid either alone or in combination mirroring the most common therapeutical approach used for the treatment of tuberculosis. With respect to BCG infection, no significant differences were observed in CFU after 1 day of treatment (Figure 4c); only the free rifampicin was able to reduce the bacterial colonies. However, a significant difference was observed after 72 hours of treatment, where both rifampicin and isoniazid-encapsulated polymersomes elicited a clear reduction in bacteria compared to the free drug (Figure 4c). Notably, the rifampicin/isoniazid co-loaded polymersomes completely eradicated the intracellular BCG after 72 hours (no CFU detected). Similar results were observed with *M. tuberculosis* infected THP-1 cells (Figure 4d). In this case, after 24 hours of treatment, the multiple drug co-loaded polymersomes significantly reduced bacterial burden compared to the controls. Moreover, this drug formulation was also able to eradicate intracellular *M. tuberculosis* after 72 hours of treatment (Figure 4d).

**Figure 4:**
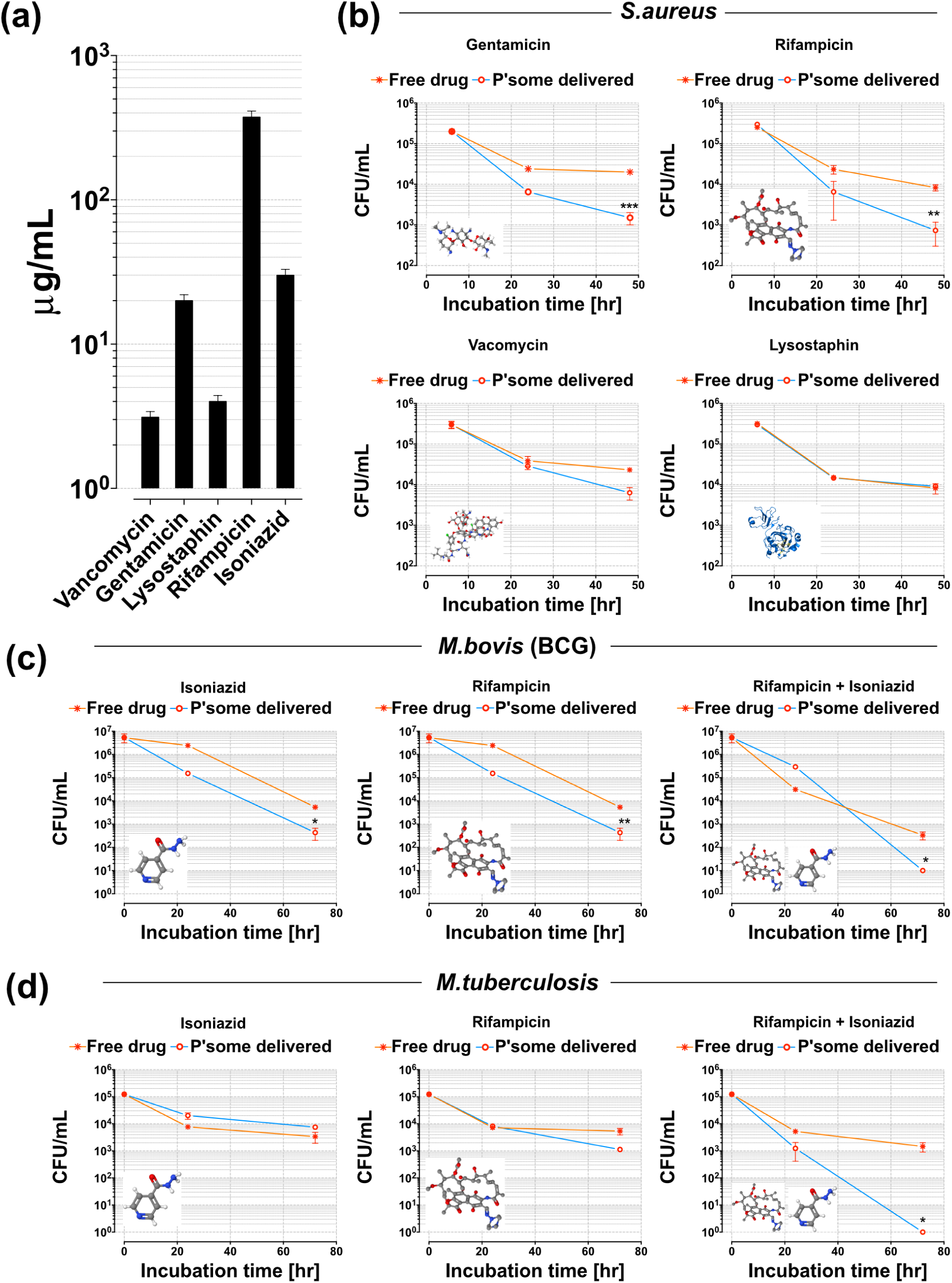
Polymersome mediated drug delivery in human macrophages. **(a)** Concentration of drug encapsulated within PMPC-PDPA polymersomes measured by HPLC. **(b)**THP-1 macrophages infected with *S. aureus* (M.O.I of 5:1) for 6 hours. Following infection, gentamicin was added to the media to kill extracellular bacteria. Macrophages were subsequently treated with polymersomes encapsulating gentamicin, rifampicin, vancomycin, or lysostaphin. At 6, 22, and 46 hours macrophages were lysed and plated on a BHI agar plate for bacterial colonies to be counted (One way ANOVA *\*\*p* < 0.01, *\*\*\*p* < 0.001, error bars = SEM, n=3). Viability (CFU) analyses of BCG **(c)**, and **(d)** *M. tuberculosis* after 24 and 72 hours of incubation with the different formulations (One way ANOVA with *\*p* < 0.05 and *\*\*p* < 0.01).

Following the *in vitro* characterisation, we moved to a relevant *in vivo* model, choosing the *Danio rerio* (zebrafish) embryo. These animals are optically transparent, allowing observation of polymersome targeting and delivery over time in the same animal. The availability of fluorescent transgenic lines labelling immune cell populations allows imaging of macrophages and neutrophils.^30^ Furthermore, there are well-established zebrafish models of human-relevant infections of *S*. aureus^31,32^ and *M. marinum* (a close relative of human TB complex, and a natural pathogen of fish species).^33,34^ We first tested the ability of PMPC-PDPA polymersomes to target macrophages using a zebrafish embryo with macrophages expressing mCherry protein and neutrophils expressing GFP protein (Figure 5a). Polymersomes co-localise effectively with macrophages, but not neutrophils, confirming mononuclear phagocytes targeting also in zebrafish. Most importantly, confocal fluorescence microscopy demonstrated PMPC-PDPA polymersomes internalise within cells infected by *S. aureus* (Figure 5b), with high level of co-localisation with the intracellular pathogen. Fluorescently labelled lysostaphin delivered by polymersomes into infected phagocytes co-localised well with intracellular *S. aureus* (Figure 5c). We corroborated these data in a tuberculosis model, by infecting zebrafish embryos with *M. marinum* and followed by microinjection with rhodamine-labelled polymersomes (**Figures 5d-g**). Ten minutes following injection (the shortest time possible to mount the samples onto the microscope slide), zebrafish macrophages infected with *M. marinum* expressing mCrimson (cyan) have already taken up the polymersomes, which co-localise with the pathogen intracellularly (Figure 5d), whilst the polymersome fluorescence is still detectable in the circulation (white arrows). Similarly, after 30 minutes, polymersomes were located freely in the circulation and within macrophages containing intracellular bacteria (Figure 5e). At 1 day post polymersome injection (d.p.i.), there was a diminished fluorescence signal in the blood-stream while macrophage levels remained high, indicating that the majority of polymersomes had been taken-up (Figure 5f). Interestingly, polymersomes continued to be evident within macrophages at 3 days post injection (Figure 5g). This is an important observation, as granulomas (a hallmark of both *M. marinum* and human TB infection) begin to form at this stage. The development of this granuloma shares similar pathology to the human tuberculous granuloma. These data provide an important insight into the *in vivo* targeting of the intracellular bacterial niche in macrophages by polymersomes to allow local release of a drug. Within this niche, microorganisms are able to evade the immune response and avoid killing by antibiotics delivered by traditional means. The majority of commercially available antibacterial drugs possess poor intracellular pharmacokinetics, which can limit their ability to treat intracellular infections.

**Figure 5:**
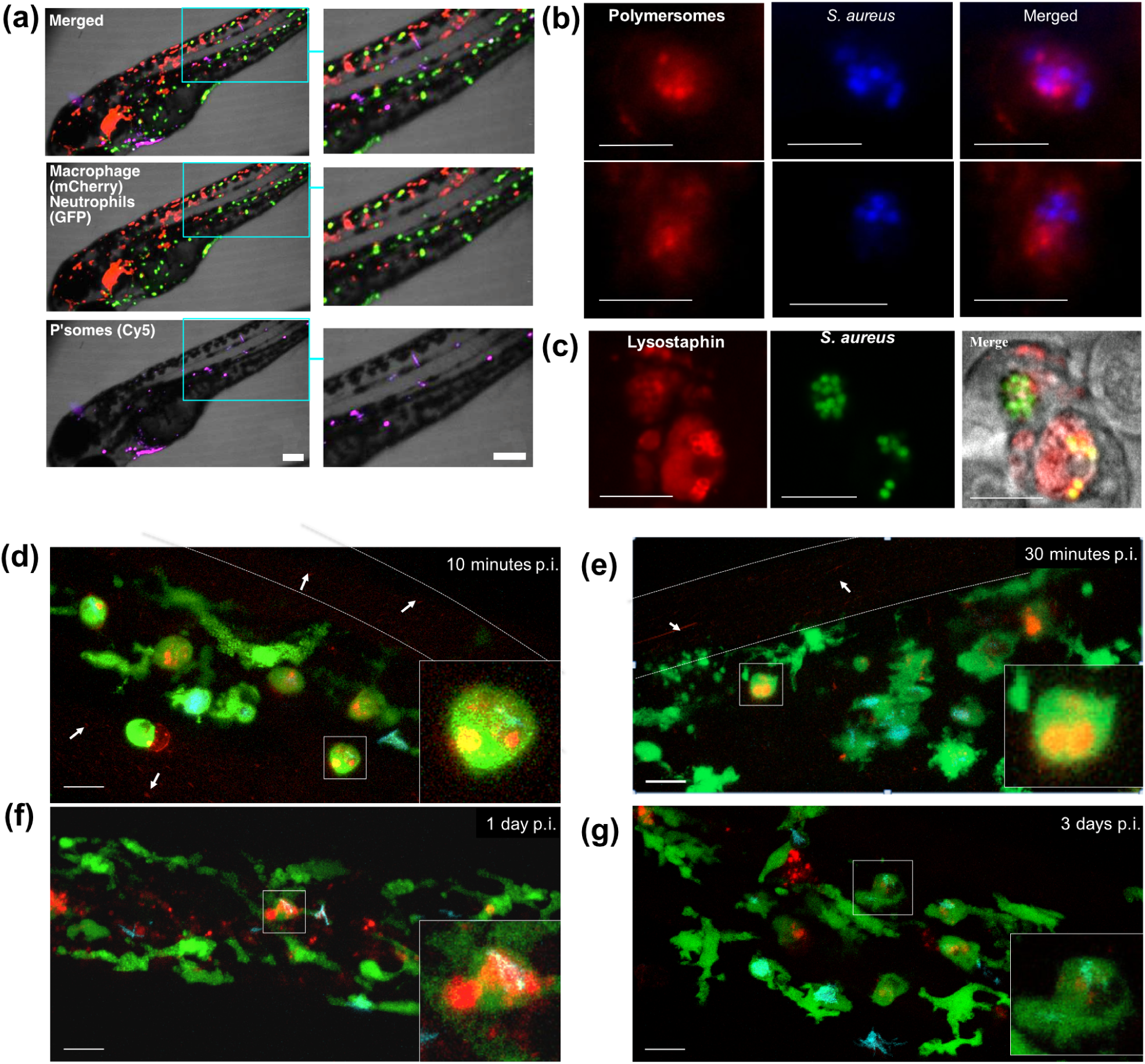
Polymersomes in zebrafish embryos infection models. **(a)** Cy5 labelled polymersomes (magenta) were injected into zebrafish embryos whose neutrophils are modified to expressed GFP (green) and macrophage to express mCherry (red). The embryos were imaged 2hr after the injection confirming strong correlation between the polymersomes and macrophages. Scale bar = 200 *μ*m.**(b)** Cy5-labelled polymersomes (red) were injected in embryos infected with *S.aureus* expressing a CFP (blue). Both bacteria and polymersomes were found in the embryo macrophages with a high co-localisation between the two. **(c)** Cy3-labelled lysostaphin encapsulated within polymersomes and injected in zebrafishes infected with GFP expressing *S.aureus*. Both the drug model and the bacteria were found to colocalise within the embryo macrophage. Confocal imaging of zebrafish embryos with GFP expressing macrophages infected with *M. marinum* expressing mCrimson (cyan), and inoculated with rhodamine-labelled polymersomes (red). Uptake imaging 10 minutes **(d)** and 30 minutes **(e)** after injection. Polymersomes are still visible in the circulatory system (dashed lines). Polymersome uptake 1 day **(f)** and 3 days **(g)** post injection. Scale bar = 20 *μ*m.

To demonstrate the therapeutic impact of polymersomes in zebrafish, we tested the ability of polymersomes encapsulated antibiotics to reduce bacterial burden *in vivo*. Zebrafish embryos were infected with mCherry-expressing *M. marinum*, and with GFP expressing *S. aureus.^31^* In the *S. aureus* infection model, zebrafish received an injection of 1200 CFU, at which dose the infection is either cleared or overwhelms the fish. Zebrafish begin to succumb to the infection after approximately 40 hours post infection (h.p.i.), so this time-point was used as an output to determine the extent of zebrafish infection.^31^ To compare the effect of encapsulated antimicrobials and free antimicrobials to treat *S. aureus* infection, zebrafish embryos (2 d.p.i.) were injected with *S. aureus* followed by a second injection of drug loaded polymersomes 20 hours later. We assessed the efficacy of the four drugs tested *in vitro*, lysostaphin, vancomycin, gentamicin and rifampicin (**Figures 6a-b**). In agreement with the *in vitro* results, only encapsulated rifampicin and gentamicin treatment improved the outcome of infection. Lysostaphin and vancomycin did not change the outcome of infection, with similar numbers to the control groups showing high numbers of bacteria (Figure 6a). The polymersomes-encapsulated rifampicin was the most effective treatment, resulting in a reduction in the bacterial CFU and preventing the fish from succumbing to overwhelming infections. Polymersomes did improve considerably the output with the rifampicin formulation getting very low CFU and with survival close to 100%. The efficacy of polymersomes delivered rifampicin was confirmed using a second *in vivo* model, the *M. marinum* infected zebrafish model of TB. In this case, mCherry-expressing fluorescent bacteria were microinjected, and 24 hours later an injection of the polymersomes-encapsulated drugs (or controls) was performed. As was the case for *S. aureus* infected zebrafish, rifampicin-encapsulated polymersomes significantly reduced the *M. marinum* burden *in vivo*, compared to the same concentration of free drug (**Figures 6b**).

**Figure 6:**
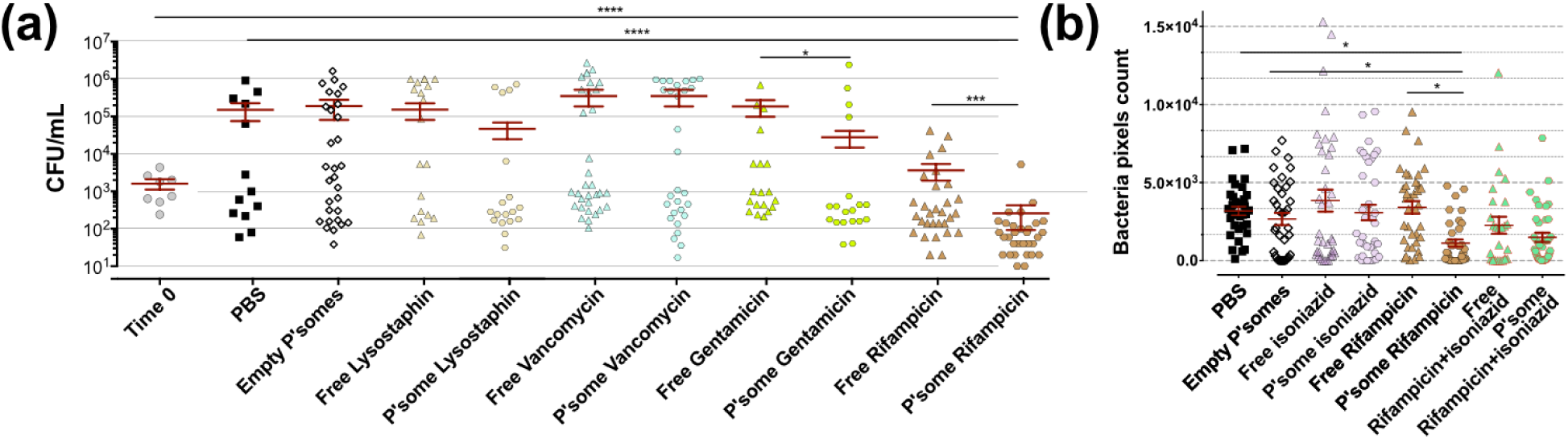
Commonly used antimicrobials encapsulated in polymersomes can be used to treat infection *in vivo*. **(a)** Zebrafish embryos 2 d.p.if were injected with *S. aureus* (data time 0) followed by a second injection 20 hours later with either PBS, empty polymersomes, free drug and polymersomes loaded with lysostaphin, vancomycin, gentamicin, and rifampicin. Zebrafish were then left for 20 hours before being homogenised and plated on BHI agar for viable colony counts. Graphs show the total number of CFU after treatment (Kruskal-Wallis test with Dunn’s multiple comparison *p <0.05, **p <0.01 and ***p <0.001). **(b)** Quantification of mCherry expressing *M. marinum* bacterial burden in zebrafish embryos treated with empty polymersomes, free drugs, and polymersomes loaded with rifampicin, isoniazid, and their combination. (ANOVA test comparison with *p < 0.05).

## Conclusions

Finding alternative and more effective solutions to bacterial infections is becoming increasingly important with the rise of antimicrobial resistant bacteria rendering many therapies ineffective. In addition, serious diseases like TB require long-term treatments, which usually need doses of a combination of antibiotics for long periods (six months), with a consequent rise of serious side effects, and bacterial resistance. New therapies, which can selectively target only infected phagocytes, are nowadays required in order to improve the efficacy while limiting off target side effects. In this work, we have demonstrated that PMPC-PDPA polymersomes are an ideal candidate for targeting mononuclear phagocytes either after i.v. or topical administration, showing tremendous potential in using this approach for those diseases where these cells are critical actors. As a proof-of-concept, we showed that PMPC-PDPA polymersomes can be loaded with a large variety of antibiotics, including proteins (lysostaphin), small peptides (vancomycin), glycols (gentamicin), poorly water-soluble organics such as quinones (rifampicin) and functionalised pyridines (Isoniazid), thus covering a large repertoire of possible chemistries. We have shown that polymersomes can deliver antibiotics to treat intracellular pathogen-related infections, and to potentially decrease the dose and duration of treatment required for bacterial eradication. Both *in vitro* in human cells and *in vivo* experiments demonstrated that these nanoscopic synthetic vesicles were specifically internalised by macrophages, without inducing toxicity, through a combination of dynamin-independent endocytosis and phagocytosis. We have demonstrated that drug-encapsulated polymersomes were able to reduce *S. aureus*, BCG, *M. tuberculosis*, and *M. marinum* bacterial burden, again using *in vitro* and *in vivo* approaches. Antimicrobial-loaded polymersomes were more effective compared with the same concentration of free drug, and in some cases were able to eradicate the intracellular microorganisms completely. We thus believe this technology can be exploited to reduce the effective dose required for therapy, with a consequent potential reduction in antimicrobial resistance. In addition, encapsulation of antimicrobials could help completely eradicate infection from the host more rapidly, by direct delivery of drug to the immune system to enhance the host-pathogen response.

## Methods

### PMPC-PDPA copolymer synthesis

In a typical ATRP procedure, a 100 mL round bottom flask equipped with a magnetic stir bar and a rubber septum was loaded with 2-methacryloyloxyethyl phosphorylcholine (MPC, 5 g, 16.9 mmol), 2-(4-morpholino)ethyl 2-bromoisobutyrate (ME-Br) initiator (189 mg, 0.7 mmol) and 6 mL ethanol, and this solution was deoxygenated by purging N2 for 1 h under stirring at r.t. Then, 2,2’-bipyridine (bpy) ligand (212 mg, 1.4 mmol) and Cu(I)Br (97 mg, 0.7 mmol) were added as solids whilst maintaining the flask under a mild positive N2 pressure. The [MPC]:[ME-Br]:[CuBr]:[bpy] relative molar ratios were 25:1:1:2. The reaction was carried out under a N2 atmosphere at 30 °C. After 90 minutes, a solution of 2-(diisopropylamino)ethyl methacrylate (DPA, 12.3 g, 57.6 mmol)in ethanol (15 mL), previously deoxygenated by purging N2 for 1 h at r.t., was injected into the flask. After 48 h, the reaction solution was opened to the air, diluted by addition of ethanol (≈200 mL) and left stirring for 1 h. The solution was then passed through a silica column to remove the copper catalyst. After this step, the filtrate was concentrated by rotary evaporation and dialysed using a 1 kDa MWCO dialysis membrane (Spectrum Labs, Netherland) against chloroform/methanol 2:1 (v/v) (2 × 500 mL), methanol (2 × 500 mL), and double-distilled water (4 × 2 L). At least 8 h passed between changes. After dialysis the copolymer was isolated by freeze-drying and characterised by 1H-NMR spectroscopy performed on an Avance III 600 spectrometer from Bruker (Billerica, USA), and gel permeation chromatography performed on a GPCMax equipped with an RI detector from Malvern Technologies (Greater Malvern, UK) with acidic water (0.25 vol% TFA in water) as solvent on a Novamax column (including guard column) from PSS Polymers (Mainz, Germany).

### Polymersomes production and characterisation

PMPC-PDPA self-assembly of polymersomes, as well as drugs encapsulation, was carried out using the thin film rehydration method. In particular,the polymers was first dissolved in a chloroform:methanol solution(2:1), containing also the antibiotics (rifampicin, isoniazid, gentamicin, and lysostaphin) at 1 mg/mL each. For the production of rhodamine-labeled polymersomes, Rhodamine 6B octadecylester (Sigma) was used (with a 5% molar ratio with the polymer). The solvent was then evaporated and the film was rehydrated with endotoxin/LPS-free Dulbecco’s PBS (Sigma) for a period of 4 weeks under vigorous stirring, in order to have a final polymer concentration of 10 mg/mL. After this period, the formed polymersomes were purified from the formed tubular structures and only spherical nanoparticles were isolated, according to sucrose-based density gradient centrifugation.^14^ This pre-purified samples were then further purified by size exclusion chromatography for isolating the antibiotics-encapsulated nano vesicles and removing the free drugs. This purified solution was then analysed by TEM, performed using a FEI Tecnai G2 Spirit electron microscope and/or a JEOL 2100 operating at 200 kV equipped with a CCD camera Orius SC2001 from Gatan. Copper grids were glow discharged and the sample was adsorbed onto the grid. The sample was then stained with 0.75wt% phosphotungstic acid (PTA) adjusted to pH 7.4 with NaOH. All the TEM analyses were carried out with dried samples. DLS analyses (for characterising the nanoparticles size distribution) were carried out using a Zetasizer Nano ZS (Malvern Ltd.) at a copolymer concentration of 0.25 mg/mL. DLS measurements were based on 12-14 runs, 10-second sub-runs. Samples were analysed at 25°C with a scattering angle of 173°and a 633 nm HeNe laser based on a material refractive index (RI) of 1.59, a dispersant refractive index of 1.330 and a viscosity of 0.89. Drugs encapsulation was measured by reverse-phase -HPLC measurements. This was performed with Dionex Ultimate 3000 instrument equipped with Variable Wavelength Detector (VWD) to analyse the UV absorption of the polymers at 220 nm and the enzymes signal at 280 nm. A gradient of H_2_O+Tryfluoroacetic acid 0.05% (TFA) (A) and MeOH+TFA 0.05% (B) from 0 min (5%B) to 30 min (100%B) was used to run the samples trough a C18 column (Phenomenex). The peak area was integrated by using Chomeleon version 6.8.

### *In vivo* (mice)tissue bio-distribution of polymersomes

Three-month-old male C57/BL6 mice were intravenously (i.v.) injected via the tail vein with 1 mg/mL rhodamine labeled-PMPC-PDPA polymersomes (*n* = 6per group). Control mice were i.v. injected with saline. The volume of solution injected was 8% of the total blood volume (TBV). TBV was calculated as 58.5 mL of blood per kg of body weight. At either 0.16, 0.5, 1, 2, 4, 6, 24 and 168 hours post-injection, the mice were terminally anaesthetised and transcardially perfused with phosphate buffered saline (PBS) 0.1 M pH 7.4. The mice were humanly terminated by cardiac puncture at pre-set time points and then perfused with PBS to remove residual blood from the organs. The plasma concentration of polymersomes was measured after centrifugation of the whole blood at time intervals (0.16, 0.5, 1, 2, 4, 6 and 24), and the relative fluorescence was measured using Xenogen IVIS 100 *in vivo* Imaging System using exposure time of 5 s and a field of view of 4×4cm. The intensity of light emission of each organ was quantified as [photons/second]/ [*μ*-icrowatt/square centimetre] and normalised by the untreated sample. To determine the interactions of Fluorescently labeled polymersomes with different types of mice blood cells B cells, T cells, monocytes, neutrophiles and RBCs, we separate the different fractions using untoched neutrophil isolation kit, Murine Peripheral Blood Neutrophil Isolation - Easysep kit and Easyplate magnet. Each organ (GI, liver, kidneys, spleen, lungs, muscles, testis, thymus, bone marrow, spinal chord and the brain were removed and the weight of each organ determined separately. The organs collected were stored at -80°C for further analysis. The amount of fluorescent signal from Rhodamine 6G (λ_*Ex*_ = 560nm, λ_*Em*_ = 600nm) in each organ was measured using Xenogen IVIS 100 *in vivo* Imaging System using exposure time of 5s and a field of view of 4×4 cm. The intensity of light emission of each organ was quantified as [photons/second]/ [μ-icrowatt/square centimetre] and normalised by the untreated sample. The radiant efficiency of the control organ was subtracted from the treated organ. The results were then reported as total radiant efficiency ([p/s] / [*μ*W/cm2]) and as percentage of the total fluorescence calculated summing all the organs measured. All procedures involving animals were approved by and conformed to the guidelines of the Institutional Animal Care Committee of The University of Sheffield. We have taken great efforts to reduce the number of animal used in these studies and also taken effort to reduce animal suffering from pain and discomfort. For the monocyte sub-population characterisations, C57/BL6 mice were maintained under specific pathogen-free conditions. All animals were fed on a standard chow pellet diet with free access to water and maintained on a 12-hour light-dark cycle. Animal work was performed in accordance with Home Office regulations Animals (Scientific Procedures) Act 1986. Mice were administered with 1 mg/mL of rhodamine labelled polymersomes I.V. and analysed as described. For the analysis of circulating monocytes, peripheral blood was collected and PBMCs enriched by Ficoll density gradient centrifugation. The PBMC fraction was stained and subsequently analysed by flow cytometry. CD11b (M1/70), Ly6C (HK1.4), and CD115 (AFS98) Abs were purchase from Biolegend, and F4/80 (CI:A3-1) from AbD Serotec. All procedures involving animals were approved by and conformed to the guidelines of the Institutional Animal Care Committee of The University of Sheffield, University College London, and University of Ghent. We have taken great efforts to reduce the number of animal used in these studies and also taken effort to reduce animal suffering from pain and discomfort.

### Cell culture and *in vitro* uptake

Human monocytes (THP-1 cell lines) were differentiated to macrophages through incubation with 5 ng/mL of phorbol 12-myristate 13-acetate (PMA, Sigma) for 48 hours on 24/96 well plates for cell viability/uptake quantification respectively, and on glass-bottom dishes (ibidi) for confocal analyses.^15^ We chose this PMA concentration as it has been found to not undesirable regulate genes expression. For cell viability, the Thiazolyl Blue Tetrazolium Blue (MTT, Sigma) method was used. Briefly, cells were seeded at a concentration of 5-10^3^ cells/well in a 96 well plate overnight (O.N.). Increasing concentrations of polymersomes were then added in the growth media, namely 0.1, 0.5, and 1 mg/mL, for periods of 24, 48, and 72 hours. The medium growth was then removed and an acidified solution of isopropanol was added to dissolve the water-insoluble MTT formazan. The solubilised blue crystals were measured colorimetrically at 570 nm (plate reader ELx800, BioTek). Viability assays were also carried out incubating cells with 10 *μ*M Acetoxymethyl (AM) Calcein staining (Invitrogen) for 1 hour, followed by confocal microscopy analyses (Leica TCS SP8). For uptake quantification, THP-1 cells were incubated with rhodamine-labeled polymersomes (0.1 mg/mL) for 8, 24, and 72 hours, followed by 3 steps of PBS washing and SDS-based cell lysis. Cell debris were then removed by centrifugation, and the rhodamine-polymers present in the surnatant quantified by HPLC.

### *In vivo* uptake in zebrafish macrophages

Adult zebrafish were maintained according to standard procedures. All experiments were performed on embryos 5 days post fertilisation (d.p.f.) or under. Transgenic strains used were the Tg(mpx:GFP)i114,^30^ and the Tg(fms:GFP)sh377.^35^ In zebrafish S. *aureus* imaging experiments, 2 d.p.f LWT zebrafish embryos were injected with 1200 CFUs of CFP-labelled S. *aureus* followed by an injection of 10 mg/mL rhodamine labelled polymersomes 1 hour later (10% Rhodamine-PMPC-PDPA, 90% PMPC-PDPA). Zebrafish were incubated for 2 hours at 28 °C before analysis by fluorescence microscopy using a Nikon TE-2000 U microscope. *M. marinum* experiments were performed using *M. marinum* strain M (ATCC #BAA-535) containing the pSMT3-mCrimson vector or pSMT3-mCherry vector.^34^ Tuberculous granuloma formation is enhanced by a mycobacterium virulence determinant. Liquid cultures were prepared from bacterial plates with 50 *μ*g/mL hygromycin as previously described.^34^ Specificity of the zebrafish host transcriptome response to acute and chronic mycobacterial infection and the role of innate and adaptive immune components.^36^ Injection inoculum was prepared from overnight liquid cultures with an OD600 of 1, after washing in PBS/Tween 80, and resuspending in 2% polyvinylpyrrolidone40 (PVP40)/PBS.^34^ Injection of *M. marinum* into zebrafish embryos was performed into the blood forming region of the caudal vein at 28-30 h.p.if.^37^ Here 200 CFU, in a volume of 1 nL, were injected. 1 d.p.i.,3 nL of Rhodamine labeled polymersomes was injected into the circulation via the Duct of Cuvier.^37^ Once injected, the embryos were mounted in 1% low melting point agarose and fluorescent confocal images and time lapses were generated using a Leica TCS SPE-II microscope using a 40x objective (water immersion, HCX PL APO, 1.10NA).

### *In vitro* and *in vivo* quantification of bacterial burden

THP-1 cells were differentiated to macrophages in a 96-well plate as previously described. Then we carried out infection with BCG, and M. tuberculosis with a Multiplicity of Infection (M.O.I.) of 10:1 for 24 hours, using antibiotics free RPMI medium. Cells were then washed 3 times in PBSt o remove the excess of microorganisms, and incubated with RPMI medium (CTRL), empty polymersomes (CTRL -), polymersomes encapsulated with Rifampicin, Isoniazid, or combination of both, and free Rifampicin, Isoniazid, or combination of both free drugs (the antibiotics were all at the same final concentration of 1 *μ*g/mL). We tested all these formulations for 24 and 72 hours, then macrophages were lysed with 0.05% SDS, and the CFUs quantified with the SPOTi assay. In particular, we carried out serial dilutions of each lysis solution, and 10 *μ*L of them were aliquoted on Middelbrock 7H11 agar medium for a period of 25-30 days, or until some colonies were visible. Colony counting was carried out manually. For THP-1 *S.aureus* experiments, mid-log S. *aureus* (Newman strain), were centrifuged at 10,000G for 1 minute and resuspended in 1 mL PBS. 106 CFUs were added to each well (MOI of 5:1). The cells were then placed on ice for 1 hour followed by a further 5 hours in a 37°C incubator (total 6 hours incubation). After incubation, gentamicin was added to the media (150 *μ*g/mL) and the cells were left for 30 minutes in an incubator to kill the extracellular bacteria. The samples were removed from the incubator, washed twice with PBS and then replaced with RPMI media containing 15 *μ*g/mL of gentamicin and the treatment or control was added. At each specified time point (6.5 hours, 22 hours, and 48 hours post infection) the media was removed, the cells were washed twice with PBS and then 250 *μ*L of 1% Saponin (Sigma Aldrich) was added to lyse the cells. The macrophages were left in the Saponin for 12 minutes in a 37°C incubator and then an additional 750 *μ*L of PBS was added to the cells and the wells were mixed thoroughly with a pipette. 10 *μ*L of the lysed cells were taken and diluted in a 96 well plate with six 1/10 serial dilutions. Three 10 *μ*L drops from each dilution were placed onto a labelled blood agar plate, incubated overnight at 37°C and the number of viable colonies were counted. For zebrafish *in vivo S. aureus* experiments, 2 d.p.if. LWT zebrafish were injected with 1200 CFUs of GFP-labelled S. *aureus*. 18 hours after injection zebrafish were viewed under a fluorescent dissecting microscope (Leica MZ10F) and zebrafish with visible abscesses were discarded. 20 hours post infection, zebrafish were injected with 0.5 nL of 1mg/mL polymersomes with 37.5 *μ*g/mL of encapsulated rifampicin or their subsequent controls. Zebrafish embryos were incubated at 28°C for a further 20 hours following polymersome injections and were then homogenised using the PreCellys 24-Dual (Peqlab). The homogenates were serially diluted onto BHI agar plates, placed in a 37°C room and the number of viable colonies was manually counted the following morning. In order to quantify the *in vivo* microorganism burden, injection of *M. marinum* into zebrafish embryos was performed into the blood forming region of the caudal vein at 28-30 h.p.f. 100 CFU, in a volume of 3 nL, were injected. 1 d.p.i., 1 nL of PBS (Ctrl), empty polymersomes, free Rifampicin (3.6 mM) or Rifampicin-encapsulated polymersomes (3.6 mM) were injected into the circulation via the Duct of Cuvier. Embryos were imaged at 4 d.p.i. on a wide field Leica DMi8 using a 2.5x objective (air, HC FL PLAN, 0.07NA) with images generated with a Hamamatsu Orca Flash 4.0 V2 camera. Bacterial burden was analysed suing pixel counting software as previously described.^34^ Data were analysed using one-way ANOVA (with Bonferroni post-test adjustment) in Prism 6.0 software (GraphPad Software, Sand Diego, CA, USA).

## Acknowledgements

We thank the EPSRC (EP/G062137/1) for the initial *in vivo* evaluation and ND and GB salary, ERC for the MEViC ERC-STG project for part of the consumable, GB and LR salary, the Royal Society Newton Fellowship for most TB related consumables and LR salary, The MRC doctoral training account for JR studentship, the Wellcome trust for PME fellowship, the NC3R for AP salary.

---

**Figure S1:**
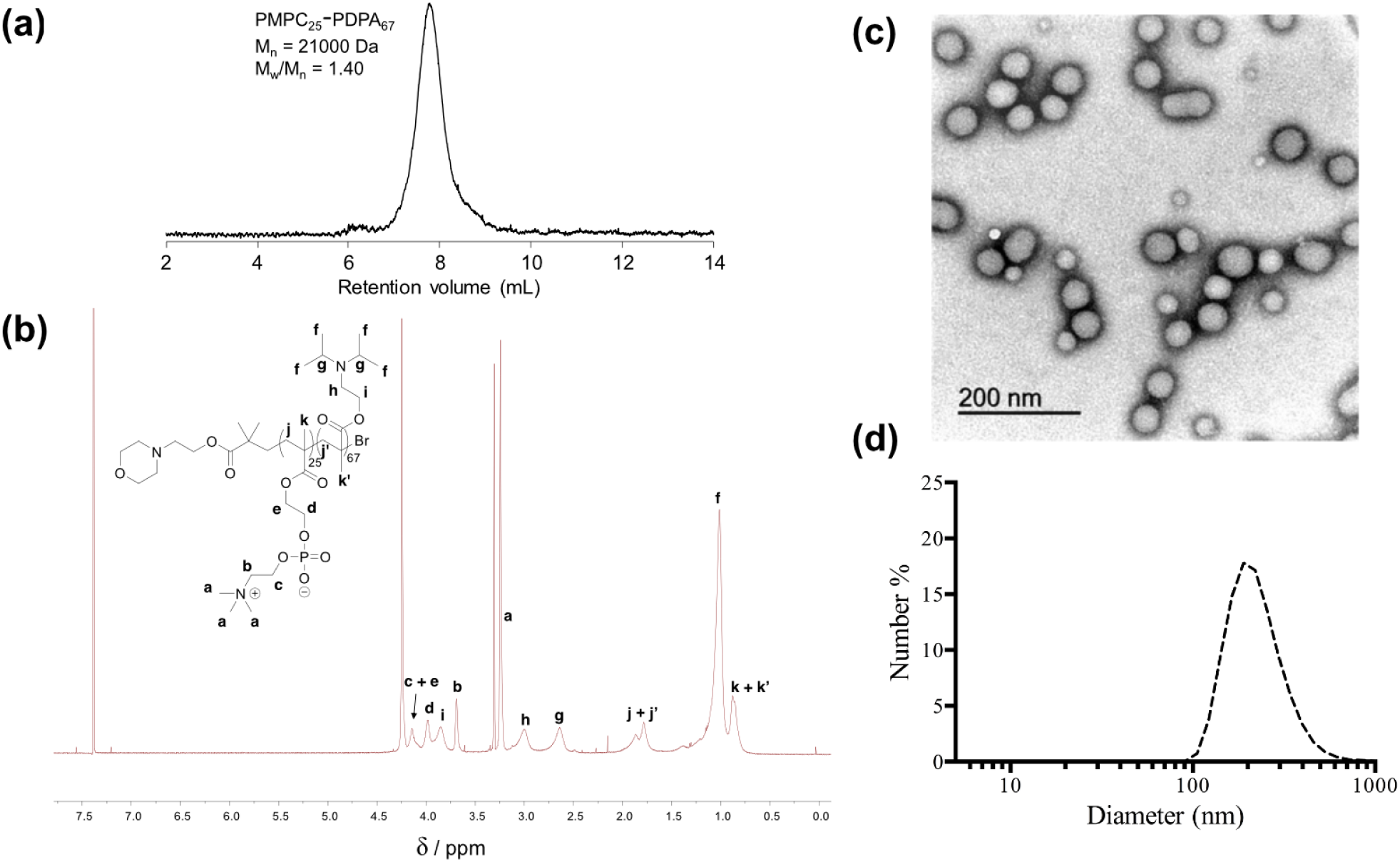
Polymer characterisation and polymersomes formation. (**a**) GPC chromatogram of PMPC25-PDPA67 analysed in DI water + 0.25%(v/v)TFA. (**b**) 1H-NMRspectrum of PMPC25-PDPA67 in CDCI3/CD3OD 3:1 (v/v). (**c**) Transmission electron micrograph of PMPC-PDPA polymersomes stained with phosphotugstic acid. (**d**) Polymersomes size distribution measured by dynamic light scattering.

**Figure S2:**
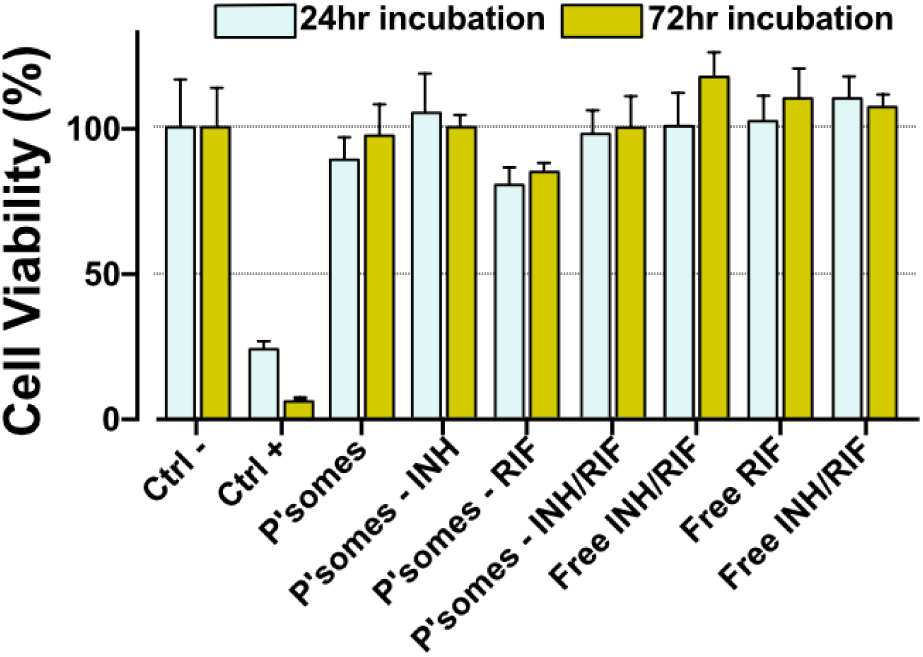
Polymersomes biocompatibility. Viability assays (MTT) of THP-1 cells incubated with (unloaded) polymersomes, and with antibiotic-loaded (rifampicin, isoniazid, and combination of both) polymersomes. Ctrl-: Cells treated with PBS; Ctrl+; DMSO 5%; [P’some]: 1 mg/mL; [RIF]: 30 *μ*g/mL; [isoniazid]: 3 *μ*g/mL.

**Figure S3:**
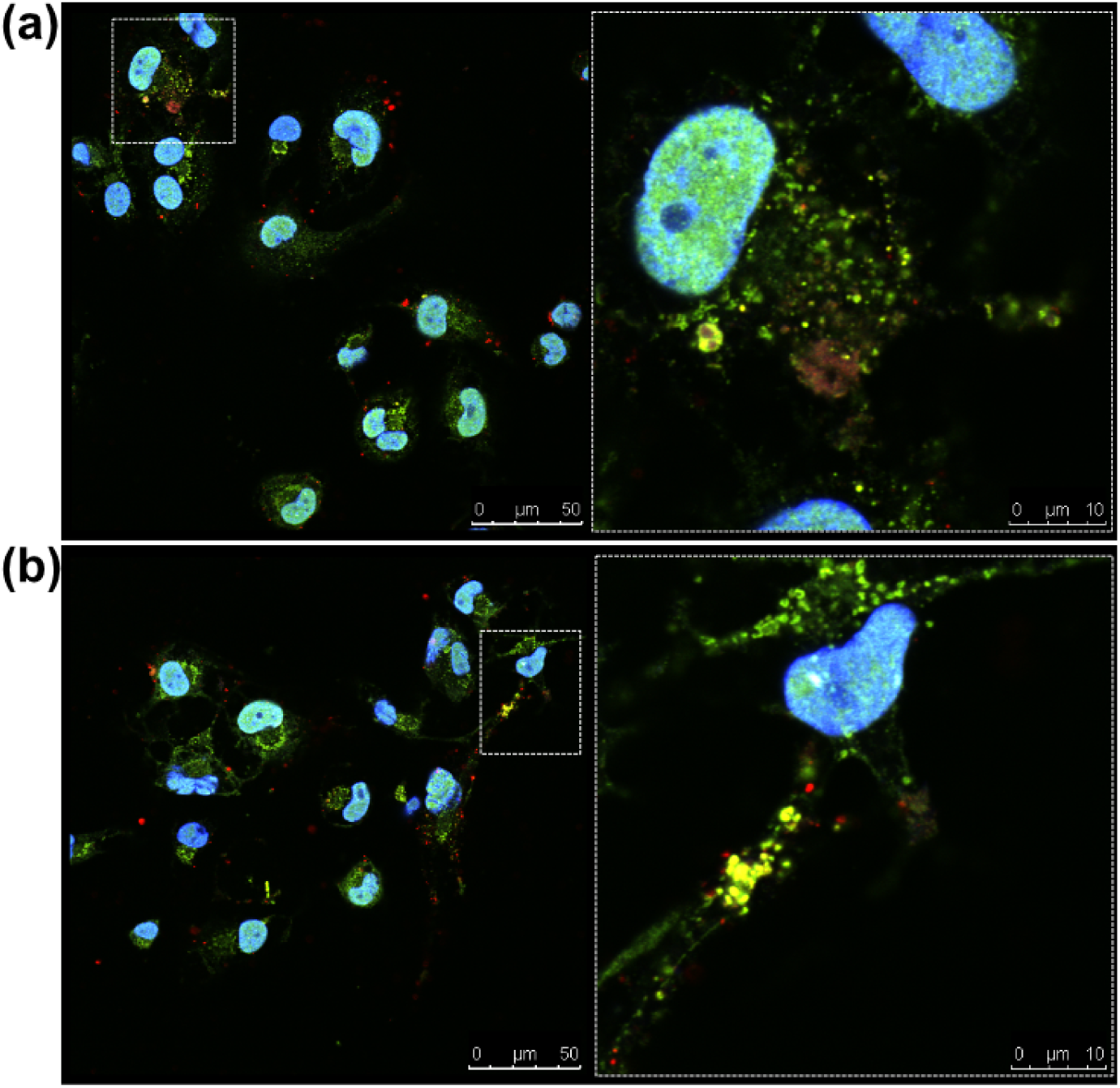
Polymersome trafficking at long incubation time. Confocal co-localisation analyses between LAMP1 (green) and Polymersomes (red) in THP-1 cells after 24h **(a)** and 72h **(b)**. Yellow signal: merge between LAMP1 and polymersomes.

**Figure S4:**
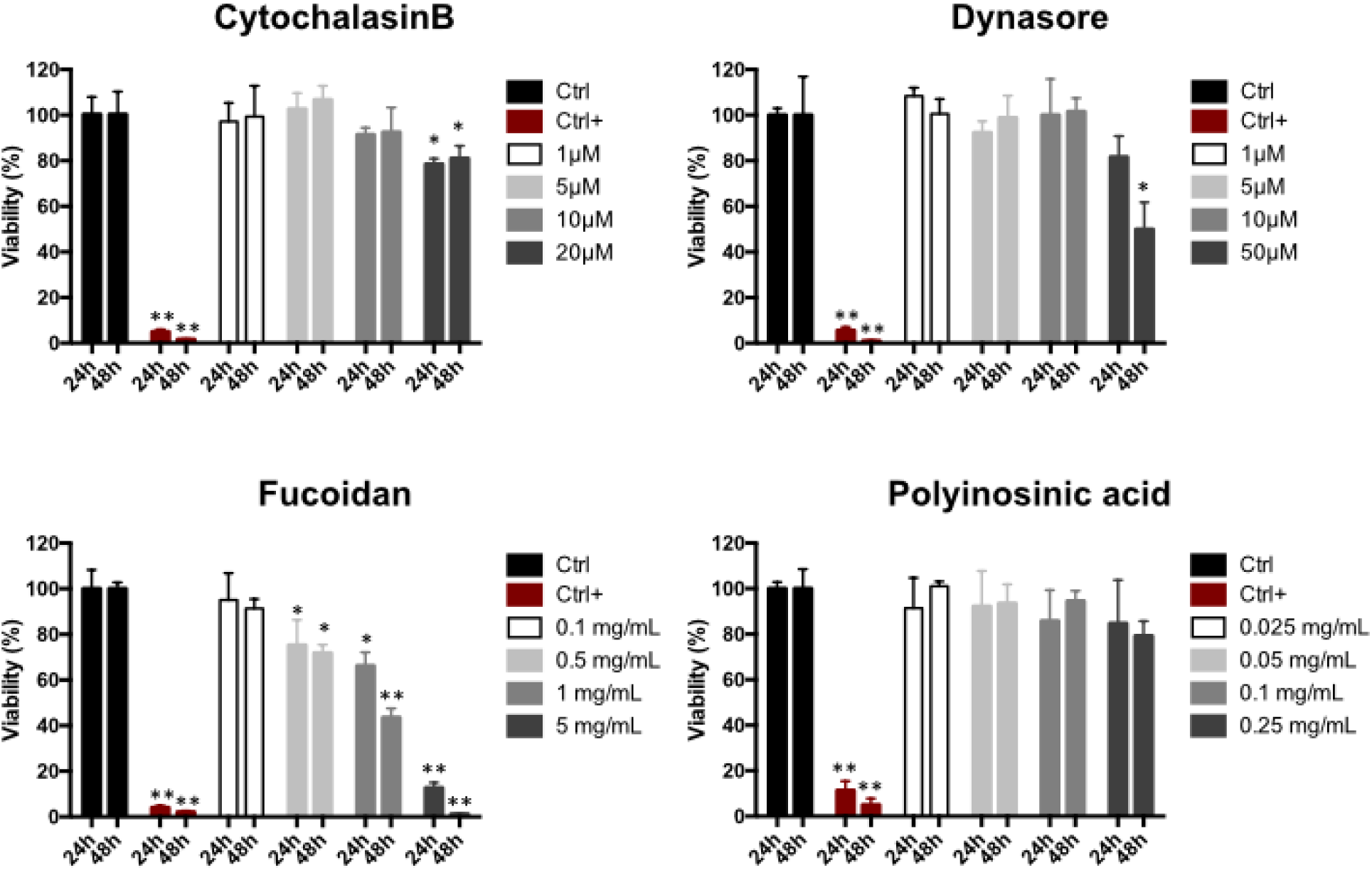
Effects of cell uptake inhibitors on Macrophages. MTT assays of THP-1 cells incubated with CytochalasinB (actin inhibitor), Dynasore (dynamin inhibitor), Fucoidan (Scavenger Receptor A and B inhibitor), and Polyinosinic acid (Scavenger Receptor A inhibitor and Toll like III receptor agonist). t-test with *p<0.05 and **p <0.01.

**Figure S5:**
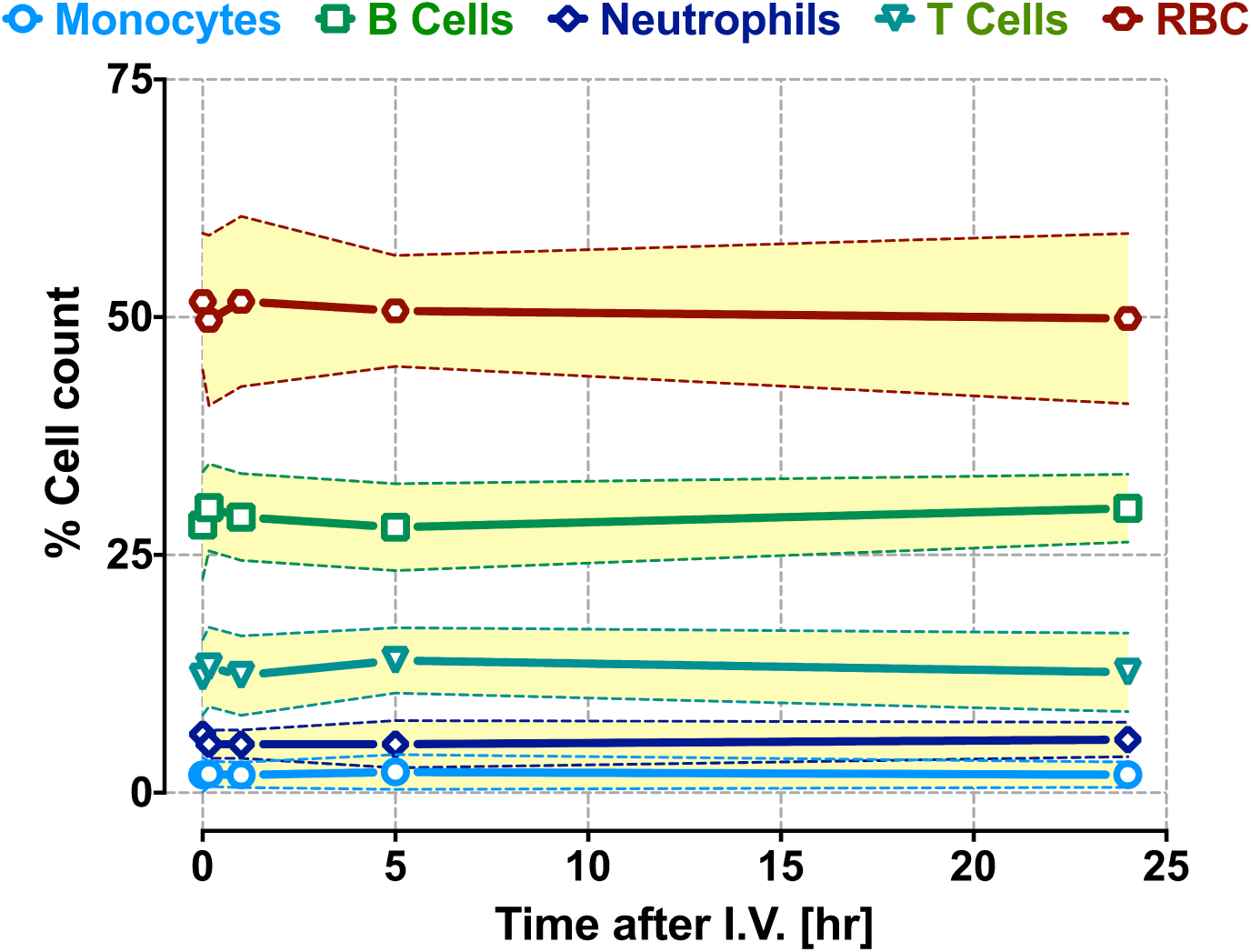
Polymersome effect on blood cells. Red and white cell count as a function of the time after polymersome IV injection

